# Diversity and some bioactivities of soil actinomycetes from southwestern China

**DOI:** 10.1101/692814

**Authors:** Yi Jiang, Guiding Li, Qinyuan Li, Kun Zhang, Longqian Jiang, Xiu Chen, Chenglin Jiang

**Author notes:** **Corresponding authors**: Yi Jiang; Chenglin Jiang.

## Abstract

With the natural medicine exploring, the actinomycetes (actinobacteria) have gotten more and more recognition. 815 soil samples were collected from six areas in the southwestern China. 7063 purified strains of actinomycetes were isolated from these samples by using four media. The 16S rRNA gene sequences of 1998 selected strains of the 7063 were determined, and the phylogenetic analysis was carried out. The diversity of actinomycetes is analyzed. Total 33 genera of actinomycetes as the purified cultivation were identified from these soil samples. 14, 13, 5, 9 and 26 genera of actinobacteria were identified from E, A, B, D, C and F area respectively, and the communities of actinomycetes are very different from each other. The diversity of Xishuangbanna (F area) is the richest, and 26 genera were isolated. That of Emei and Qingcheng Mountain (C area) is monotone, and only five genera were isolated. 158 of 1998 strains (7.9 %) are possible new species. Antimicrobial activities of 1070 selected strains against 11 bacteria and fungi were tested using agar well diffusion method, and biosynthetic genes of type I and II polyketide synthases (PKS I, PKS II), nonribosomal peptide synthase (NRPS) and polygene cytochrome P450 hydroxylase (CYP) of 1036 selected strains were detected by PCR. High rate of antimicrobial activity and the four antibiotic biosynthetic genes existed in these actinomycetes. Results of this study indicates that firstly more unknown actinomycetes can be obtained from soil samples, specially primeval tropical forests. Second, isolation methods for actinomycetes must be continually improved and improved.

**Importance:** First, discarding repeat of a mass of known microbes and compounds is very difficult. Second, pharmaceuticals development can still not overtake the increase of resistance of pathogens to antibiotics, and new diseases continuously and fleetly occur. Soil actinomycetes of these regions are researched few by microbiologist yet. In order to get much more unknown actinomycetes for discovery of new bioactive metabolites, the actinomycetes of forest soil in southwestern China were isolated and identified. Anti-microbial activities and synthetic gene clusters’ of four kinds of antibiotics of some selected strains were detected.

## INTRODUCTION

Actinomycetes (Actinobacteria) has been paid a great attention owing to their production of various natural drugs and other bioactive metabolites including antibiotics, enzyme inhibitors and enzymes, and has special ecological function such as fixation of air nitrogen and degradation of difficult disaggrega-tire substance in the nature world for maintaining a balance of ecosystem. Over 22,000 bioactive secondary metabolites (including antibiotics) were published in the scientific and patent literature, and about 11000 of 22000 metabolites were produced by actinomycetes by the end of 2002. About 150 antibiotics have being applied in human therapy and agriculture now. In the 150 antibiotics, bacteria produced 10-20, actinomycetes produced 100-120, and fungi produced 30-35 (Berdy 2005). There are two difficult problems of natural pharmaceutical development from microorganisms. First, discarding repeat of a mass of known microbes and compounds is very difficult. So pharmaceutical development needs very long time, tremendous investment, and manpower and material resources. Second, pharmaceuticals development can still not overtake the increase of resistance of pathogens to antibiotics, and new diseases continuously and fleetly occur. But 90 % to 99 % of microorganisms in the nature world are not cultured yet based on the research results of molecular technology (Chiao 2004; Hughes et al. 2001; Joseph et al. 2003; Pachter 2007; Zengler et al. 2002). Making the uncultured microorganisms to cultured microorganisms is one hope for getting new leader compounds for development of new natural drug. Southwestern China including Yunnan, Guizhou, Sichuan and Tibet is one of regions of the richest biodiversity in China. Soil actinomycetes of these regions are researched few by microbiologist yet. In order to get much more unknown actinomycetes for discovery of new bioactive metabolites, the actinomycetes of forest soil in southwestern China were isolated and identified. Anti-microbial activities and synthetic gene clusters’ of four kinds of antibiotics of some selected strains were detected. Some results are reported here.

## MATERIALS AND METHODS

### Collection and pretreatment of soil samples

280 soil samples were collected from primeval subtropical every-green broadleaf forest in Zhangjiajie of Hunan and Fanjing Mountain in Guizhou, China. The two sampling spots belong to Wuling Cordillera (A sampling area). 50 soil samples were from primeval subtropical every-green broadleaf forest of Huangjing in Gulin (B area), south of Sichuan. 100 samples from secondary every-green broadleaf forest in Emei and Qingcheng Mountains (C area), the western brim of Sicuan Basin. 50 samples from primeval alpine taiga of Jiuzhaigou (D area), in the north of Sichuan. 220 samples from primeval forests on various altitudes of Grand Shangri-La (E area) in the common boundary of Sichuan, Yunnan and Tibet. 85 soil samples were collected from primeval tropical rainy forest of Xishuangbanna in the south of Yunnan (Figure 1). Every soil sample was collected from 3 to 5 holes with 10 to 20 cm depth and incorporated as one sample, and put in one sterile plastic bag. Total 815 soil samples were preserving at 4 ºC before test. The soil samples were dried in room temperature for 7 to 10 days, then pretreatment at 80 ºC for 1 hour.

**Figure 1.**
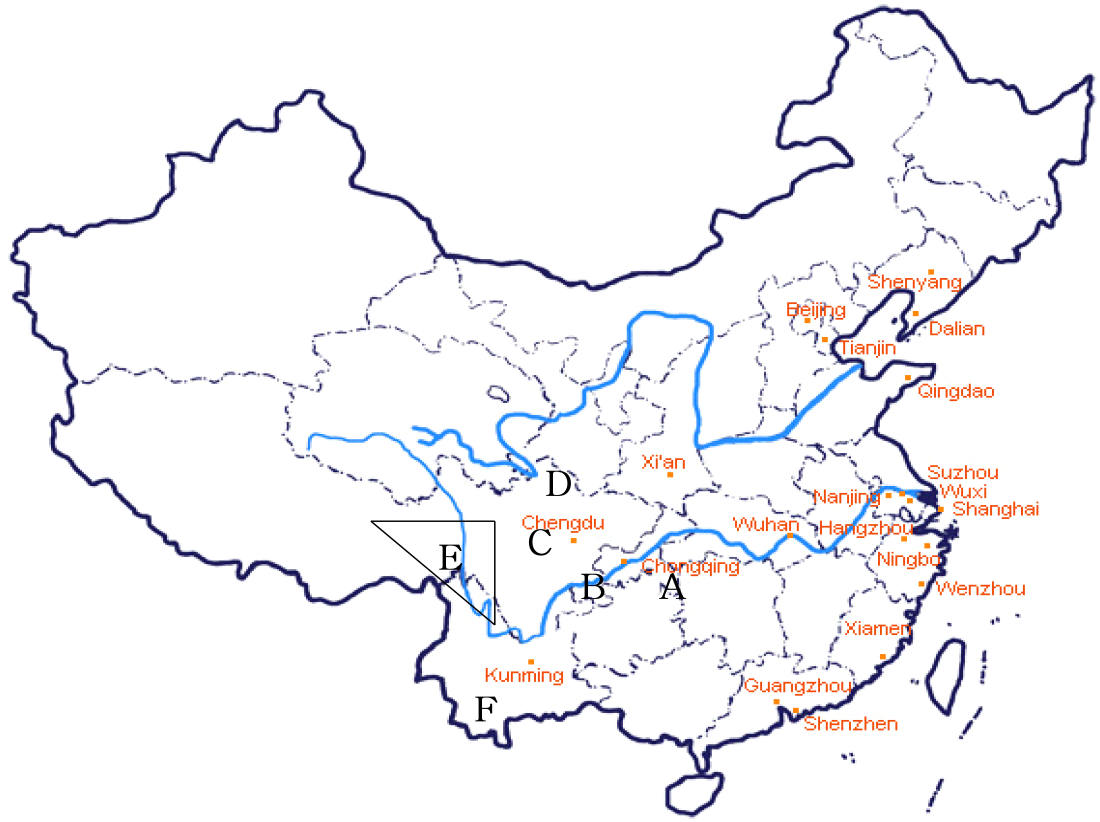
Position of sampling areas in China

### Isolation medium and method of actinomycetes

Plate dilution method was used for isolation of actinomycetes. Four media were used as following: YIM 7 (HV medium) (Hayakawa and Nonomura 1987); YIM 171 [Improved Glycerol-asparagine medium: Glycerol 10 g, asparagine 1 g, K_2_HPO_4_⋅H_2_O 1 g, MgSO_4_⋅7H_2_O 0.5 g, CaCO_3_ 0.3 g, Vitamin mixture (Hayakawa and Nonomura 1987) 3.7 mg, trace salts (Shirling and Gottlieb 1966) 1 ml, potassium dichromate (K_2_Cr_2_O_7_) 50mg, agar 15 g, distilled water 1000 ml, pH 7.7]; YIM 37 (Jiang and Xu 1997) (Improved Histidine-raffinose medium: Histidine 1 g, raffinose 10 g, Na_2_HPO_4_ 0.5 g, KCl 1.7 g, CaCO_3_ 0.02 g, MgSO_4_⋅4H_2_O 0.05 g, FeSO_4_⋅7H_2_O 0.01 g; Vitamin mixtures 3.7 mg, cycloheximide 50 mg, nystatin 50 mg, potassium dichromate 50 mg, agar 15 g, distilled water 1000 ml, pH 7.2); YIM 212 (Jiang et al. 2006) (Mycose-proline medium: Mycose 5 g, proline 1 g, (NH_4_)_2_SO_4_ 1 g, NaCl 1 g, CaCl_2_ 2 g, K_2_HPO_4_ 1 g, MgSO_4_⋅7H_2_O 1 g, vitamin mixtures 3.7 mg; potassium dichromate 50 mg, agar 15 g, distilled water 1000 ml, pH 7.2).

### Characterization of actinomycetes

Cultivation of cell, extraction of DNA, PCR and sequencing of 16S rRNA gene and phylogenetic analysis of 1998 actinomycete strains were carried out with the methods described in previously paper (Jiang et al. 2008). The strains were characterized at a genus and species level.

### Determination of antimicrobial activity

920 actinomycete strains were fermented in YIM 61 broth (Soybean meal 20 g, glucose 10 g, peptone 4 g, K_2_HPO_4_ 1 g, MgSO_4_⋅7H_2_O 0.5 g, NaCl 1 g, CaCO_3_ 2 g, distilled water 1000 ml, pH 7.8) on shaker at 28 ºC for 7 days. The fermented broth was used for determining whether inhibition against four bacteria and 7 pathogenic fungi of crop.

### Detection of synthetic gene cluster of four antibiotics

Extraction of DNA of 1036 strains was carried out by using the methods described by Xu et al (Xu et al. 2003). Genes of type I and II polypeptides synthetase (PKS I, PKS II), nonribosomal peptide synthetase (NRPS) and polygene cytochrome P450 hydroxylase (CYP) were detected with the methods of references (Ayuso-Sacido and Genilloud 2005; Metsä-Ketelä et al. 1999; Hwang et al. 2007) respectively.

## RESULTS

### Isolation effect of four media

Total 7063 actinomycete strains were isolated as purified culture from the 815 samples collected from before-mentioned six sampling areas, Wuling Cordillera (A), Huangjing (B), Emei and Qingcheng Mountains (C), Jiuzhaigou (D), Grand Shangri-La (E) and Xishuangbanna by using four media (Table 1). 1790 strains including 46% of Streptomycetes and 54 % of non-streptomycetes were isolated with HV (Hayakawa and Nonomura 1987). The rates of streptomycetes and non-streptomycetes were 64 % and 36 % by using improved Glycerol-asparagine medium (YIM 171), and 58 % and 42 % by improved Histidine-raffinose medium (YIM 37) respectively. 1384 strains with 41 % of streptomycetes and 59 % of non-streptomycetes were isolated by Mycose-proline medium (YIM 212). These results indicate that YIM 171 and YIM 37 can be applied for isolation of streptomycetes. YIM 212 and HV can be used for isolation of non-streptomycetes actinomycetes. Potassium dichromate is an effective selective inhibitor of fungi and bacteria for selective isolation of rare actinomycetes. No fungi and few bacteria grew on all of agar plates containing 50 mg/L of potassium dichromate during incubation.

**Table 1.** Amount of isolated actinomycete strains from soil samples collected from five areas in southwestern China with four media

| Sampling area | YIM 171 |  | YIM 37 |  | YIM 7 (HV) |  | YIM 212 |  |
| --- | --- | --- | --- | --- | --- | --- | --- | --- |
|  | Streptomycete | Other | Streptomycete | Other | Streptomycete | Other | Streptomycete | Other |
| A* | 221 | 108 | 155 | 115 | 87 | 112 | 148 | 167 |
| B | 187 | 78 | 26 | 30 | 168 | 273 |  |  |
| C | 458 | 258 | 70 | 59 | 128 | 73 |  |  |
| D | 120 | 106 | 55 | 73 | 43 | 85 |  |  |
| E | 293 | 145 | 305 | 195 | 299 | 336 | 355 | 514 |
| F | 358 | 203 | 112 | 48 | 105 | 81 | 61 | 139 |
| Total | 1637 | 898 | 723 | 520 | 830 | 960 | 564 | 820 |
| % | 64 | 36 | 58 | 42 | 46 | 54 | 41 | 59 |

\*A=Wuling Cordillera; B=Huangjing; C=Emei and Qingcheng Mountains; D=Jiuzhaigou; E=Grand Shangri-La; F=Xishuangbanna

### Composition of actinomycetes in Wuling Cordillera (A Sampling area)

Wuling Cordillera is situated in a common boundary of Chongqing, Hubei, Hunan and Guizhou. The cordillera is from east to west, area is about 100,000 km^2^, and the physiognomy belongs to calcareous. Fanjing Mountain, one of two sampling spots is situated in the northeast of Guizhou, and belongs to primeval subtropical every-green broadleaf forest. Altitude for sampling area is 1150 m to 2493 m. Zhangjiajie, another sampling spot, is a national park and primeval subtropical every-green broadleaf forest in the northwest of Hunan. Altitude of sampling area is 350 m to 1360 m. Main representative plants are *Castanopsis*, *Quercus, Machilu*, *Cunninghamia, Cinnamomum* and *Ginlgo* (Hou 2001). 1113 strains of actinomycetes were isolated from 280 samples collected from Wuling Cordillera with four media. 432 of the 1113 strains were selected for identification. A part of sequences (700 bp~1100 bp) of 16S rRNA gene of these strains were determined. Phylogenetic trees were constructed based on the 16S rRNA gene sequences. The results showed that the 432 strains belonged to 6 suborders, 8 families and 14 genera, *Streptomyces*, *Micromonospora*, *Dactylosporangium*, *Catellatospora*, *Sphaerosporangium*, *Streptosporangum*, *Actinomadura*, *Nonomuraea*, *Nocardia*, *Rhodococcus*, *Arthrobacter*, *Microbacterium*, *Mycobacterium* and *Pseudonocadia*. Figure 2 is showing the phylogenetic tree based on 16S rRNA gene sequences of some strains and related species of the family *Streptosporangiaceae* (data of rest families not shown). *Sphaerisporangium* is described recently by Ara and Kudo (2007). YIM 48771^T^ and YIM 48782^T^ are two new species of the genus *Sphaerisporangium* based on polyphasic taxonomy, and was published (Cao et al. 2009). YIM 48783, YIM 48778 and YIM 48775 were possible new species of *Streptosporangium*, YIM 48789 was possible new species of *Nocardia*, and YIM 48790 was possible new species of *Dactylosporangium* (data unshown).

**Figure 2.**
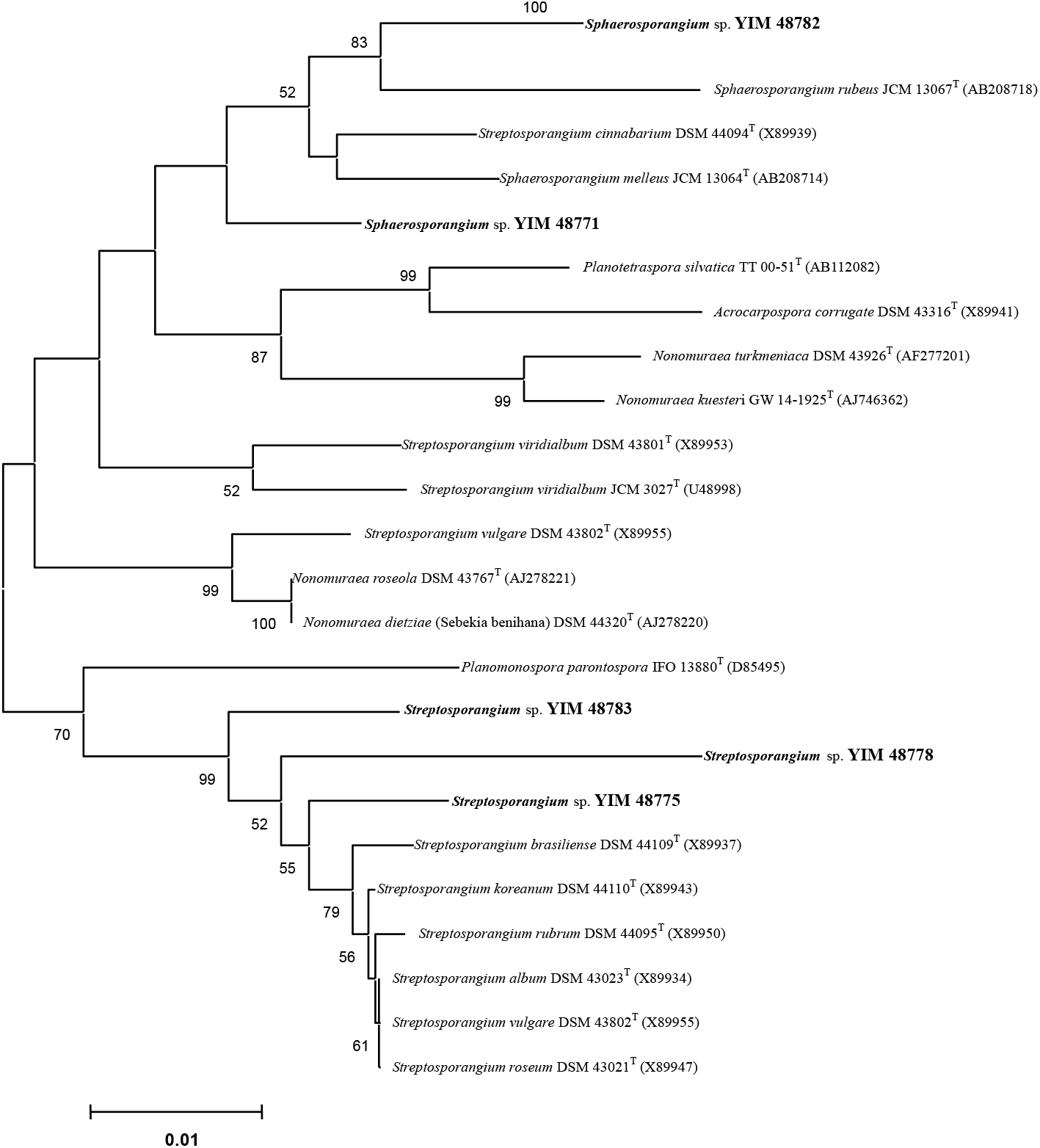
Phylogenetic tree based on 16S rRNA gene sequences of cultured strains from Wuling Mountain and related species of the family *Streptosporangiaceae.* Bar: 1% sequence divergence

### Composition of actinomycetes in Huangjing (B area)

Huangjing is in Gulin, a common boundary of Sichuan, Guizhou and Yunnan, at north latitude 28°, altitude 720 m to 1720 m, and about 3182 km^2^. The vegetation belongs to primeval subtropical every-green broadleaf forest. Main representative plants are *Quercus gilliana, Castanopsis platyacantha, Cinnamomum camphora., Phoebe* sp.*, Pinus armanddii, Liriodenfron chinesie, Fokienia* sp.*, Rhodoleia* sp.*, Machilus ichangensis, Sinarundinaria nitida* and *Cyathea spinulosa* (Hou 2001). 762 strains of actinomycetes were isolated from 50 soil samples collected in this area. 242 of the 762 strains were identified by phylogenetic analysis based on 16S rRNA sequences, and consisted of 7 suborders, 8 families and 13 genera, *Mycobacterium, Nocardia, Rhodococcus, Micromonospora, Pseudonocardia, Saccharomonospora, Actinomadura, Nonomurae, Promicromonospora, Nocardioides, Verrucosispora, Actinopolymorpha* and *Streptomyces.* Figure 3 is phylogenetic tree of the 13 genera.

**Figure 3.**
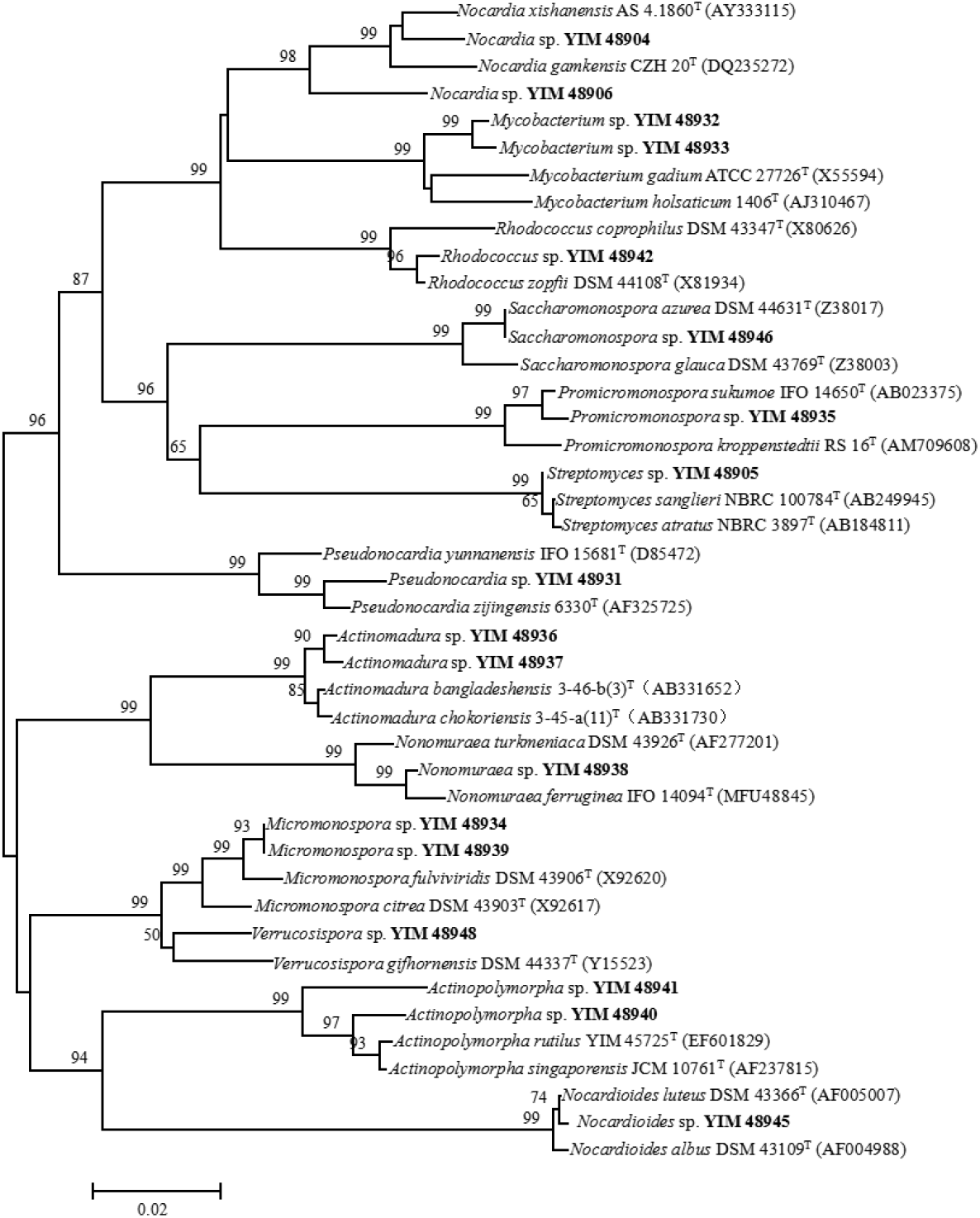
Phylogenetic tree based on 16S rRNA gene sequences for the some cultured strains of the 13 genera from Huangjing (B) and related species. Bar: 2% sequence divergence

### Composition of actinomycetes in Emei and Qingcheng Mountains (C area)

Emei Mountain, one of two spots of C area, is situated in the western brim of Sichuan Basin, at north latitude 29°, altitude 520 m to 3080 m, and about 154 km^2^ range. The vegetation belongs to secondary subtropical every-green broadleaf forest. Main representative higher plants are *Davidia involucrata*, *Cyathea spinulos, Castanopsis platryacantha, Abies ceratacantha, Machilus ichangensis*, etc. Qingcheng Mountain, another sampling spot, is situated in the northwestern brim of Sicuan Basin, at north latitude 31°, altitude 570 m to 1120 m. The vegetation in there belongs to secondary every-green broadleaf forest. Representative plants are Genera *Quercus*, *Castanopsis, Lithocarpus, Cyclobalanopsis, Pinus, Cunninghamia, Juniperus, Sabina, Alnus* and *Sinarundinaria* (Hou 2001). 1046 strains were isolated from 100 soil samples collected from this area. 125 strains of them were identified by phylogenetic analysis (Figure 4) based on 16S rRNA gene sequences, and consisted of 4 suborders, 4 families and 5 genera, *Mycobacterium, Nocardia, Promicromonospora, Dactylosporangium* and *Streptomyces.* 1 strain belonged to other bacteria, *Paenibacillus*.

**Figure 4.**
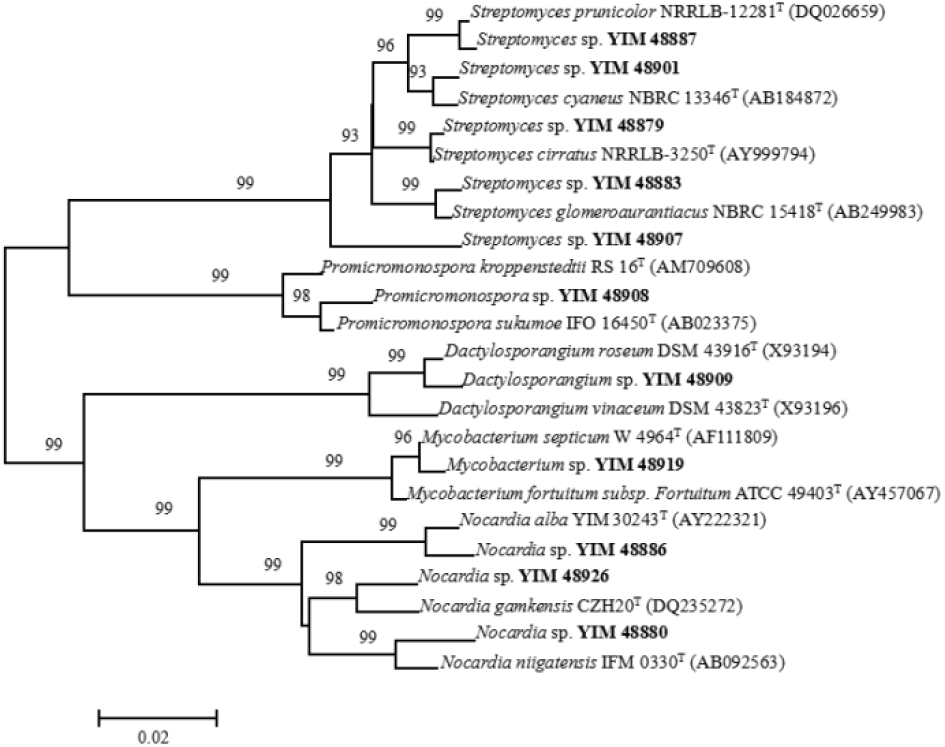
Phylogenetic tree based on 16S rRNA gene sequences for a part of cultured strains of the 5 genera from Mountains Emei and Qingcheng (C), and related species. Bar: 2% sequence divergence

### Composition of actinomycetes in Jiuzhaigou (D area)

Jiuzhaigou is one of famous beauty spots in China, situated in the north of Sichuan, at north latitude 34°, altitude 1500 m to 3100 m. The vegetation in there has been protected very well, and belongs to primeval alpine taiga. Main plants are genera *Picea, Abies, Larix, Salix, Quercus, Rhododendron* and *Sinarundinaria* (Hou 2001). 482 strains were isolated from 50 samples collected in this area. 209 strains of them were identified by phylogenetic analysis based on 16S rRNA sequences. They were consist of 6 suborders, 7 families and 8 genera, *Mycobacterium, Promicromonospora, Kribbella, Micromonospora, Actinomadura, Nonomuraea, Pseudonocardia* and *Streptomyces* (Figure 5).

**Figure 5.**
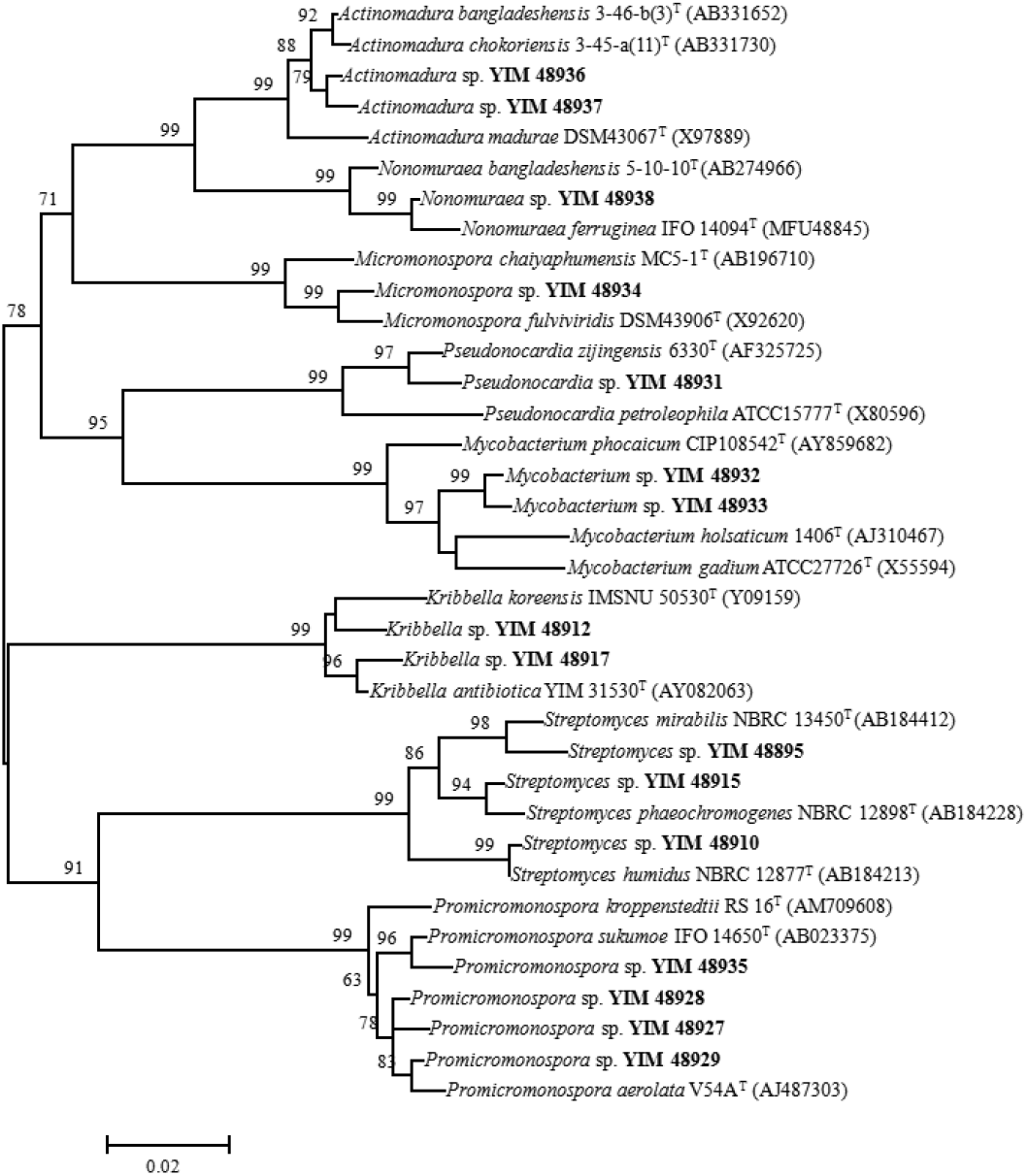
Phylogenetic tree based on 16S rRNA gene sequences for a part of cultured strains of the 8 genera from Jiuzhaigou (D) and related species. Bar: 2% sequence divergence

### Composition of actinomycetes in Grand Shangri-La (E area)

Grand Shangri-La belongs to Hengduan Cordillera between east longitude 94° to 102° and north latitude 26° to 34°. Snow Mountains are consists of Meili, Baimang, Haba, Yulong, Gongga, Nianbaoyuze and Nanjiabawa Snow Mountain. Lu, Lancang, Jinsha, Yalong, Dadu and Min Rivers are going through these big cordilleras. Altitude difference is over 6000 m, from the bottom of river to top of mountain. That is one of pure land, unfrequented and having many engaging stories. The highest 5257 m, the lowest 1950 m, and over 3000 m in the most of sampling areas. The most of soil belongs to dark brown and black clay (alpine meadow soil), and humus is over 25 cm thickness. Grand Shangri-La is one of the richest biodiversity areas in China. Forest in there has been protected very well. The vegetation belongs to every-green broadleaf forest from 1950 m to 3000 m, and main plants are *Castanopsis delavayi*, *Lithocarpus craibianus*, *L. Pachyphyllus*, *Schima argentea*, *Pinus yunnanensis* and *Alnus nepalensis;* alpine taiga and boskage from 3000 m to 4000 m, and distributing *Quercus pannosa*, *Rhododendrom rubiginosum*, *Sinarundinaria* sp., *Iris bulleyyana* and *Clinelymus nutans*; alpine meadow from 4000 m to 5250 m, and distributing *Juniperus wallichiana*, *Queercus guayavaefolia*, *Primula serrantifolia*, *Kobresia stiebritzian, Eremopogon delavayi*, *Festuca ovina*, *Salix calyculata*, *Rhododendron roxieanum*, *Rh. Traillianum* (Hou 2001). 220 soil samples were collected in the region.

Table 2 showed the amount of isolated actinomycetes in different altitude by using four media. 1463 strains were isolated from 80 soil samples collected from 1950 m to 3000 m high, and average 18 strains for one soil sample. 697 strains were isolated from 80 samples of 3000 m to 4000 m, and 9 strains for one sample. 282 strains from 60 samples of 4000 m to 5250 m, and 5 strains for one sample. Total 2442 strains were isolated from the 220 soil samples. The amount of isolated actinomycetes is decreasing along with raise of altitude. These results are similar with that in Ailao Mountain, Yunnan, China (Jiang and Xu 1996).

**Table 2.**
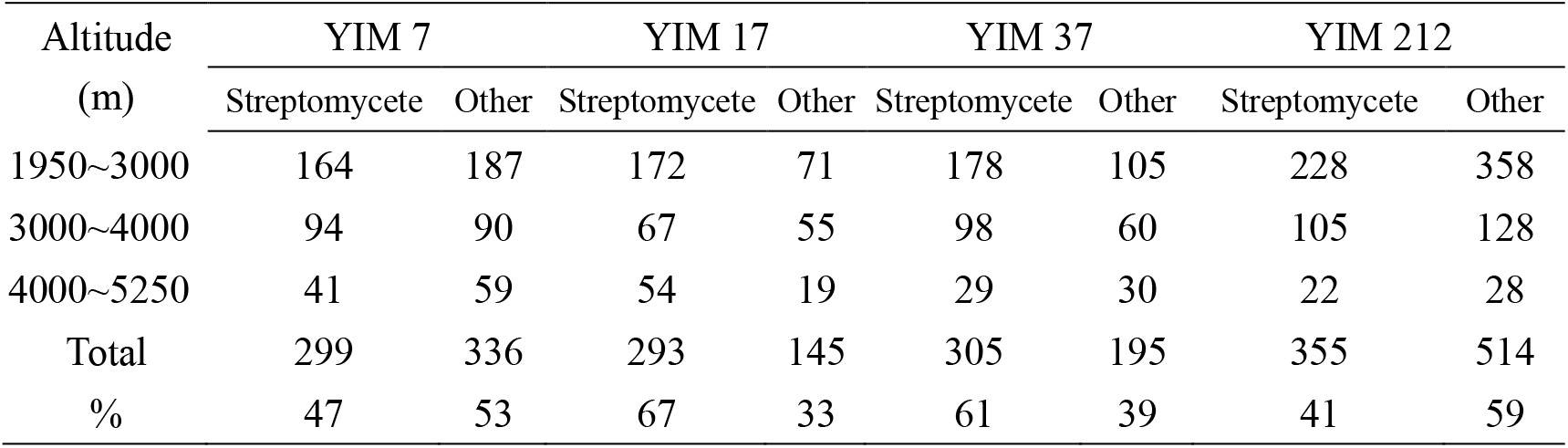
Amount of isolated actinomycete strains of soil samples collected from different altitude in Grand Shangri-La with four media

| Altitude<br>(m) | YIM 7 |  | YIM 17 |  | YIM 37 |  | YIM 212 |  |
| --- | --- | --- | --- | --- | --- | --- | --- | --- |
|  | Streptomycete | Other | Streptomycete | Other | Streptomycete | Other | Streptomycete | Other |
| 1950~3000 | 164 | 187 | 172 | 71 | 178 | 105 | 228 | 358 |
| 3000~4000 | 94 | 90 | 67 | 55 | 98 | 60 | 105 | 128 |
| 4000~5250 | 41 | 59 | 54 | 19 | 29 | 30 | 22 | 28 |
| Total | 299 | 336 | 293 | 145 | 305 | 195 | 355 | 514 |
| % | 47 | 53 | 67 | 33 | 61 | 39 | 41 | 59 |

Total 2442 strains of actinomycetes were isolated from 220 soil samples collected from E area. 436 strains were selected from the 2442 strains for identification. Sequences (700 bp~900 bp) of 16S rRNA gene of the 436 strains were analyzed. Phylogenetic tree was constructed based on the sequences. The results indicate that total 7 suborders, 13 families and 20 genera at least were isolated as pure culture. 23 strains belonged to *Nocarioides* and *Actinopolymorpha* of *Nocardioidaceae* of suborder *Propionibacterineae* (data unshown), 87 strains to *Kocuria* and *Arthrobacter* of *Micrococcaceae, Agromyces* and *Mycetocola* of *Microbacteriacea, Oerskovia* of *Cellulomonadaceae, Promicromonospora* of *Promicromonosporaceae* of suborder *Micrococcineae* (Fig. 6), 78 strains to *Nocardia* and *Rhodococcus* of *Nocardiaceae* and *Tsukamurella* of *Tsukamurellaceae* of suborder *Corynebacterineae* (data unshown), 44 strains to *Pseudonocardia* of *Pseudonocardiaceae* and *Lentzea* of *Actinosynnemataceae* of suborder *Pseudonocardineae*, 66 strains to *Dactylosporangium, Micromonospora* and *Planosporangium* of *Micromonosporineae* of suborder *Micromonosporneae* (data unshown), 42 strains to *Actinomadura* of *Thermomonosporaceae* and *Streptosporangium and Sphaerisporangium* of *Streptosporangiacea* of suborder *Streptosporangineae*, and 96 strains to *Streptomyces*. 16S rRNA gene similarities of 38 of the 436 strains with the closest type species were lower than 98.65 %. In the other word, most of them are possible new species. YIM 48868 is a new possible species of genus *Actinopolyporpha*, and was published in IJSEM (data unshown). YIM 48875 was new possible species of genus *Planosporangium* (Wiese et al. 2008). Rest 5 strains belonged other bacteria, *Methylobacterium, Massilia, Ralstonia, Roseomonas* and gram positive bacteria, *Bacillus*. Figure 6 as an example is showing the phylogenetic tree belonging to the sub-order *Micrococcineae* and related species.

**Figure 6.**
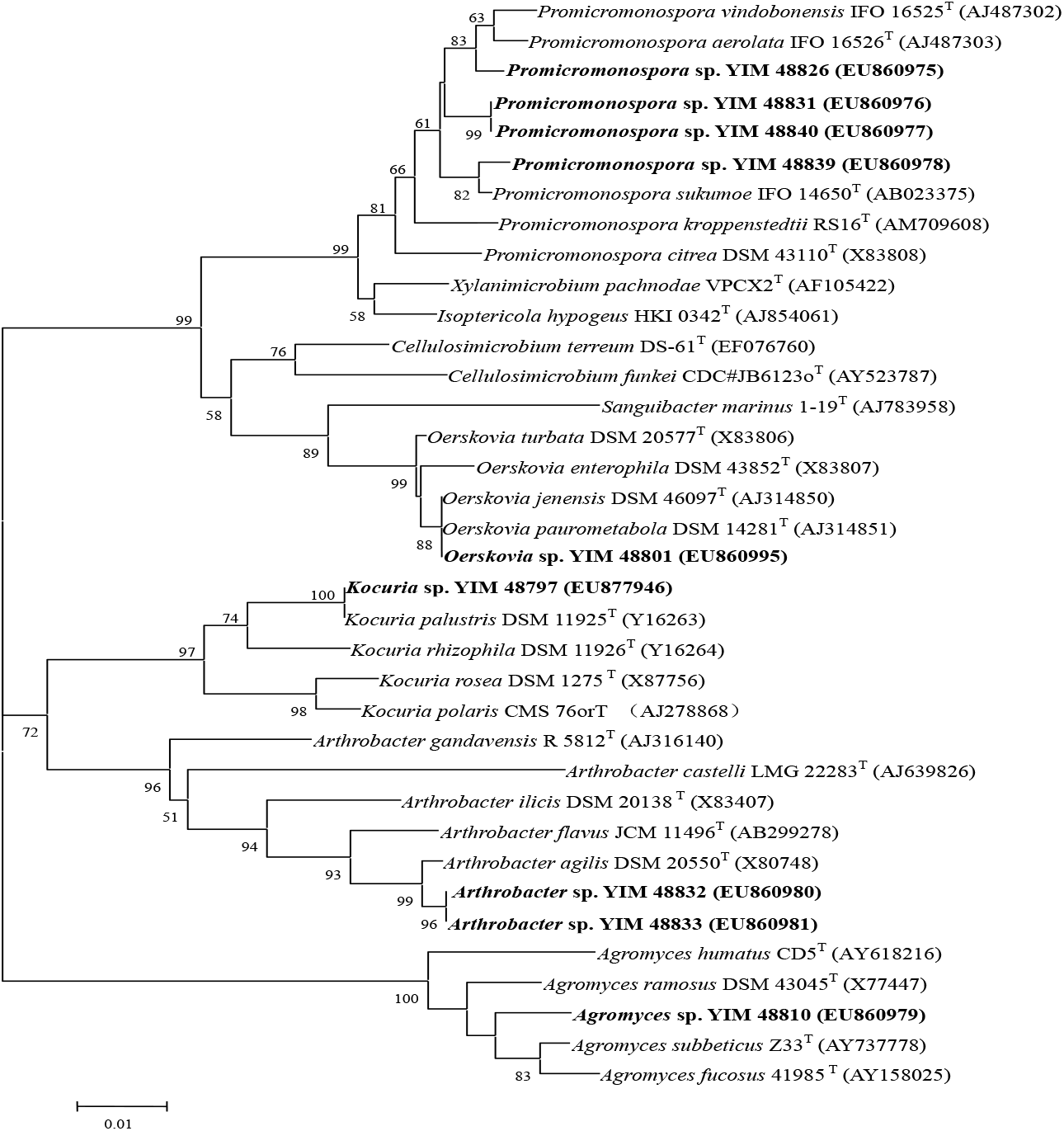
Phylogenetic tree showing the relationships among reference strains and experimental strains of sub-order *Micrococcucineae* based on 16S rRNA sequences based on neighbor-joining analyses of 1000 resampled data sets. Bar: 1% sequence divergence. Numbers in parentheses represent the sequences accession number in GenBank.

### Psychrophilic actinomycetes in E Area

Actinomycetes of 30 soil samples collected over 3700 m in E area were isolated with YIM 171 medium at 4 ºC and 8 ºC respectively. 38 of 43 strains isolated at 4 ºC were *Streptomyces*, 3 strains of *Micromonospora*, 2 of *Pseudonocardia*. 52 of 68 strains isolated at 8 ºC were *Streptomyces*, 6 of *Pseudonocardia, 5* strains of *Micromonospora*, 3 of *Nocardia* and 2 of *Actinomadura*. The results indicate that streptomycetes are still preponderant under low temperature environments (Table 3).

**TABLE 3.** Amount of isolated actinomycetes at 4 ºC and 8 ºC

| Genus | Isolation at 4 °C | Isolation at 8 °C | Total No. |
| --- | --- | --- | --- |
| *Streptomyces* | 38 | 52 | 90 |
| *Micromonospora* | 3 | 5 | 8 |
| *Actinomadura* | 0 | 2 | 2 |
| *Pseudonocardia* | 2 | 6 | 8 |
| *Nocardia* | 0 | 3 | 3 |
| Total | 43 | 68 | 111 |

All of the 111 strains can grow at 4 ºC. 97 strains of them grown well at 10 ºC to 28 ºC. 15 of the 97 can grow at 37 ºC belonging to normal temperature microbes, and 82 strains did not grow at 37 ºC belonging to psychrotolerant microbes. Rest 14 strains, including 12 *streptomyces* and 2 *Nocardia*, grown well at 10 ºC to 14 ºC, did not grow at 28 ºC, and belonged to psychrophilic actinomycetes.

### Composition of actinomycetes in Xishuangbanna (F area)

Xishuangbanna (F Area) is located in the south of Yunnan province. The region has a typical monsoon climate with a mean annual temperature ranging between 15.1◦C and 21.7◦C, and precipitation between 1200 and 2500 mm. Xishuangbanna is a region with the richest biodiversity in China. 85 soil samples were collected from tropical rain forest in Xishuangbanna national nature reserve. 1107 strains of actinomycetes were isolated from the 85 samples. 443 strains of them were identified by phylogenetic analysis based on 16S rRNA sequences. These strains belonged to 27 genera of actinobacteria, *Actinomadura, Actinoplanes, Actinopolymorpha, Agrococcus, Agromyces, Arthrobacter, Citricoccus, Dactylosporangium Friedmanniella, Jiangella, Kribbella, Lentzea, Microbacterium, Micromonospora, Mycobacterium, Nocardia, Nocarioides, Nonomuraea, Oerskovia, Planosporangium, Promicromonospora, Pseudonocardia, Rhodococcus, Saccharopolyspora, Sphaerisporangium, Streptomyces, Streptosporangium*, the community of actinomycetes was richest in this study, and members of *Actinoplanes, Agrococcus, Citricoccus, Friedmanniella, Jiangella* and *Saccharopolyspora* did not isolated in other areas in this study.

### Antimicrobial activities

Antimicrobial activities of 1070 selected actinomycete strains against 2 Gram negative and 2 positive bacteria and 7 pathogenic fungi of crop were determined. Antimicrobial spectrum of the actinomycetes from six sampling areas were different each other (Table 4). 13 and 12 of 182 strains from A area had inhibition against *Phytophthora nicotianae* and *Bacillus megaterium* respectively. 6 strains had inhibition against *Fusarium* sp. YIM 48776 (*Micromonospora* sp.) and YIM 48878 (*Streptosporangium* sp.) had high inhibition against 3 species of bacteria. YIM 48794 (*Actinomadura* sp.) had high inhibition against almost pathogenic fungi. Average 3.0% of the 182 strains from A area had antimicrobial activities against one or several of 11 test microbes. 11 of 158 strains from B area had inhibition against *Bacillus megaterium.* 8 strains had high activities against 1 to 3 species of bacteria and 1 to 4 species of pathogenic fungi at the same time. All of the 158 strains did not inhibit *Colletotrichum* sp. 6, 4 and 4 of 103 strains from C area had inhibition against *Bacillus subtills, Fusarium* sp. and *Aspergillus niger* respectively. 9 strains had high antimicrobial activities against 1 to 3 species of bacteria and 1 to 5 species of pathogenic fungi at the same time. 18, 16 and 15 of 126 strains from D area had inhibition against *Bacillus megaterium Staphylococcus aureus* and *Bacillus subtills* respectively. 12 strains had high inhibition against 1 to 3 species of bacteria and 1 to 7 species of pathogenic fungi at the same time. YIM 48915 (*Streptomyces* sp.) had high inhibition activities against 2 species of bacteria and 7 species of pathogenic fungi. YIM 120296 (*Streptomyces* sp.) had high inhibition activities against 2 species of bacteria and 7 species of pathogenic fungi. 12, 11 and 11 of 236 strains from E area had inhibition against *Bacillus subtills, Bacillus megaterium* and *Alternaria alternata* respectively. 2 strains of *Streptomyces* had inhibition against 3 species of bacteria and 4 speciea of pathogenic fungi. No strains can inhibit *Protomyces macrosporus.* 22, 19 and 14 of 224 strains from F area had inhibition against *Bacillus subtills, Bacillus megaterium* and *Staphylococcus aureus* respectively. Average 4.9 % strains from F area had antimicrobial activity, and the positive % was the highest. Total 6.6% and 6.1% of 1070 tested strains had inhibition against *Bacillus megaterium* and *Bacillus subtills* respectively, and only 1.8%, 1.4% and1.3% strains had inhibition against pathogenic fungi, *Protomyces macrosporus, Gonmatopyricularia amomi* and *Colletotrichum* sp. respectively. Average 3.5 % of the 1070 strains had antimicrobial activities against one or several tested microbes.

**TABLE 4.** Number of positive strains with Antimicrobial activities

| Antimicrobial activities | A area | B area | C area | D area | E area | F area | Total | % |
| --- | --- | --- | --- | --- | --- | --- | --- | --- |
| Tested number | 182 | 158 | 103 | 226 | 177 | 224 | 1070 |  |
| Number of positive strains: |  |  |  |  |  |  |  |  |
| *Escherichia coli* | 3 | 2 | 3 | 6 | 2 | 11 | 27 | 2.5 |
| *Staphylococcus aureus* | 9 | 8 | 3 | 16 | 6 | 14 | 56 | 5.2 |
| *Bacillus subtilis* | 3 | 7 | 6 | 15 | 12 | 22 | 65 | 6.1 |
| *Bacillus megaterium* | 12 | 11 | 2 | 18 | 11 | 19 | 71 | 6.6 |
| *Phytophthora nicotianae* | 13 | 8 | 2 | 8 | 4 | 9 | 44 | 4.1 |
| *Alternaria alternata* | 3 | 7 | 3 | 8 | 11 | 12 | 44 | 4.1 |
| *Fusarium* sp. | 6 | 5 | 4 | 8 | 2 | 11 | 34 | 3.2 |
| *Colletotrichum* sp. | 0 | 0 | 1 | 3 | 7 | 3 | 14 | 1.3 |
| *Gonmatopyricularia amomi* | 4 | 3 | 1 | 4 | 2 | 1 | 15 | 1.4 |
| *Protomyces macrosporus* | 2 | 4 | 2 | 3 | 0 | 8 | 19 | 1.8 |
| *Aspergillus niger* | 5 | 5 | 4 | 5 | 2 | 10 | 31 | 2.9 |
| Total | 60 | 60 | 31 | 94 | 59 | 120 | 461 |  |
| Average % of positive strains | 3.0 | 3.5 | 2.7 | 3.7 | 3.0 | 4.9 |  | 3.5 |

### Biosynthetic gene clusters of four kinds of antibiotics

Detection results for biosynthetic gene clusters of four kinds of antibiotics of 1036 selected strains are shown in Table 5. The results indicate that positive rate of actinomycete strains producing each gene from six sampling areas was different each other. 13, 17 and 18 of 180 strains from A area produced PKS I, PKS II and NRPS genes respectively. 8, 11 and 12 of 121 strains from B area produced the three genes respectively. 8, 7 and 5 of 92 strains from C area produced the three genes respectively. 13, 9, and 6 of 138 strains from D area produced the three respectively. 18, 6, 11 and 13 of 216 strains from E area produced PKS I, PKS II, NRPS and CYP genes respectively. 21, 22, 37 and 27 of 289 strains from F area produced the four genes respectively. These results indicate, actinomycetes from tropical rain forests are more likely to produce antibiotics. In addition, 7.8%, 7.1%, 8.6% and 5.6% of 1036 tested strains produced PKS I, PKS II, NRPS and CYP biosynthetic gene cluster respectively. Actinomycetes contained the biosynthetic gene clusters for a variety of different antibiotics, and remain one of the main sources for the discovery of new drug leads. It is also proved that actinomycetes release antibiotics to maintain the balance of soil ecosystem in nature.

**TABLE 5.** Number of positive strains with biosynthetic genes of four metabolites

| Synthesis genes of metabolites | A area | B area | C area | D area | E area | F area | Total | % |
| --- | --- | --- | --- | --- | --- | --- | --- | --- |
| Number of test | 180 | 121 | 92 | 138 | 216 | 289 | 1036 | 100 |
| Number of positive strains: |  |  |  |  |  |  |  |  |
| PKS I | 13 | 8 | 8 | 13 | 18 | 21 | 81 | 7.8 |
| PKS II | 17 | 11 | 7 | 9 | 6 | 22 | 74 | 7.1 |
| NRPS | 18 | 12 | 5 | 6 | 11 | 37 | 89 | 8.6 |
| CYP | 4 | 6 | 4 | 4 | 13 | 27 | 58 | 5.6 |
| Total | 52 | 37 | 24 | 22 | 48 | 107 |  |  |
| Average % of positive strains | 7.1 | 7.6 | 6.5 | 3.3 | 5.6 | 8.7 |  | 7.3 |

## DISCUSSION

Total 33 genera of actinomycetes as the purified cultures were isolated and identified from total 815 soil samples collected in the southwest of China, 14, 13, 5, 8, 19 and 27 genera were isolated from A, B, C, D, E and F area respectively, and the compositions of actinomycetes are very different each other. Actinomycete diversity in primeval tropical rainforest soil in Xishuangbanna (F Area) is the richest in this study. Secondly Grand Shangri-La (E area) belonging to primeval every-green broadleaf forest, and 19 genera were identified. That of Emei and Qingcheng Mountain belonging to secondary subtropical every-green broadleaf forest, the community of actinomycetes was monotone, and only five genera were isolated. Streptomycetes are common genus for all of areas. Members of *Actinomadura, Micromonospora, Mycobacterium, Nocardia*, *Promicromonospora*, and *Pseudonocardia* were isolated from soil samples of five areas. Twelve genera, *Dactylosporangium, Nonomuraea* and *Rhodococcus* were isolated from four areas. *Actinoplans, Agrococcus, Catellatospoa, Citricoccus, Friemanniella, Jiangella, Kokuria, Mycetocola, Saccharomonospora, Saccharopolyspora, Tsukamurella* and *Verrucosispora* were isolated only from one area, they were rare actinomycetes in soil (Table 6).

**TABLE 6.** Comparison of actinomycete community in different soil

|  | A* | B | C | D | E | F |
| --- | --- | --- | --- | --- | --- | --- |
| *Actinomadura* | + | + |  | + | + | + |
| *Actinoplanes* |  |  |  |  |  | + |
| *Actinopolymorpha* |  | + |  |  | + | + |
| *Agrococcus* |  |  |  |  |  | + |
| *Agromyces* |  |  |  |  | + | + |
| *Arthrobacter* | + |  |  |  | + | + |
| *Catellatospora* | + |  |  |  |  |  |
| *Citricoccus* |  |  |  |  |  | + |
| *Dactylosporangium* | + |  | + |  | + | + |
| *Friedmanniella* |  |  |  |  |  | + |
| *Jiangella* |  |  |  |  |  | + |
| *Kocuria* |  |  |  |  | + |  |
| *Kribbella* |  |  |  | + |  | + |
| *Lentzea* |  |  |  |  | + | + |
| *Microbacterium* | + |  |  |  |  | + |
| *Micromonospora* | + | + |  | + | + | + |
| *Mycetocola* |  |  |  |  | + |  |
| *Mycobacterium* | + | + | + | + |  | + |
| *Nocardia* | + | + | + |  | + | + |
| *Nocarioides* |  | + |  |  | + | + |
| *Nonomuraea* | + | + |  | + |  | + |
| *Oerskovia* |  |  |  |  | + | + |
| *Planosporangium* |  |  |  |  | + | + |
| *Promicromonospora* |  | + | + | + | + | + |
| *Pseudonocardia* | + | + |  | + | + | + |
| *Rhodococcus* | + | + |  |  | + | + |
| *Saccharomonospora* |  | + |  |  |  |  |
| *Saccharopolyspora* |  |  |  |  |  | + |
| *Sphaerisporangium* | + |  |  |  | + | + |
| *Streptomyces* | + | + | + | + | + | + |
| *Streptosporangium* | + |  |  |  | + | + |
| *Tsukamurella* |  |  |  |  | + |  |
| *Verrucosipora* |  | + |  |  |  |  |
| **Total 33** | **14** | **13** | **5** | **8** | **20** | **27** |

\*A=Wuling Cordillera; B=Huangjing; C=Emei and Qingcheng Mountains; D=Jiuzhaigou; E=Grand Shangri-La; F=Xishuangbanna. +=having

These results indicate that the diversity of actinomycetes is the richest in “primeval environments”, especially primeval tropical rainforest soil in Xishuangbanna which has not been destroyed or changed by man and keeping in a natural estate once more. The forests in Grand Shangri-La have been protected very well, actinomycete diversity is very rich, and 20 genera were isolated. The forests in Emei and Qingcheng Mountains are secondary forests, and man’s molestation is high frequency, the actinomycete diversity is monotone. Similar results were got in previous study (Jiang and Xu, 1996). So it is very important to protection of primeval forest and other primeval environments for protection of biodiversity.

Average 3.0 % of strains from A area had antimicrobial activities. The % of antimicrobial strains from B, C, D, E and F areas were 3.5 %, 2.7 %, 3.7%, 3.0 % and 4.9 % respectively, and general at 2.7% to 3.7%. Xishuangbanna’s actinomycetes not only have high proportion of antimicrobial activities, but also have strong activity. Actinomycete strains from each area had strong antimicrobial activities against one or more tested microbes. In this study, 7.8%, 7.1%, 8.6% and 5.6% of 1036 tested strains produced PKS I, PKS II, NRPS and CYP biosynthesis gene-clusters respectively. Overall, results of this study indicates that about 3.5% of actinomycete strains had antimicrobial activities, and 7.3% of the strains contained more than one biosynthetic gene clusters of four kind of antibiotics.

Some authors have been used a word “uncultivable microorganisms” to define these microorganisms which did not isolate the purified culture yet for a long time (Chiao 2004; Hughes et al. 2001; Pachter 2007). But we think, all of microorganisms should be cultured in theory. So we prefer to used “cultured microorganisms” which “have been isolated as a purified culture” and “uncultured microorganisms” which “have been isolated as purified culture up to now yet”. Based on the results of genomic research, new species (specially genus) may contain new functional genes for biosynthesis of new secondary metabolites with new bioactivity (Yang et al. 2007). Therefore discovery of novel lead compounds from new genera and new species should be very tempting. One of aims of biodiversity study of actinomycetes is provide with the unknown actinomycetes for discovery of drug lead. It is indicated from the results of phylogenetic analysis of 16S rRNA gene and DNA-DNA homology that similarity of 16S rRNA gene sequence of one strain with known species is below 98.65%, possibility of the strain being new species is 80% (**Kim** et al. 2014). 16S rRNA gene sequences of 1998 strains were determined in this study. The sequence similarity of 158 strains of them with known species are below 98.6% and 7.9 % are possible new species. These possible new species containing 1 to 4 of detected biosynthetic gene clusters of four antibiotics and antimicrobial activities will be a main source for discovery of new bioactive metabolites in the future. Results of this study indicates that first more unknown actinomycetes can be obtained from soil samples collected from primeval environment, specially tropical forests. Second, isolation methods for actinomycetes must be continually improved and improved. “The study of isolation methods is always on the way”.

## ACKNOWLEGEMENTS

This research was supported by the National Basic Research Program of China (No. 2004CB719601), the National Natural Science Foundation of China (No. 30560001, 30900002, 31270001 and 31460005), the International Cooperative Program of the Ministry of Science of Technology, P. R. China (2006DFA33550), and the “Zentrum für Marine Wirkstoffe”, which is founded by the Ministerium für Wirtschaft, Wissenschaft und Verkehr des Landes Schleswig-Holstein (Germany).

## REFERENCES

Ara, I., and Kudo, T. 2007. *Sphaerosporangium* gen. nov., a new member of the family Streptosporangiaceae, with descriptions of three new species as *Sphaerosporangium melleum* sp. nov., *Sphaerosporangium rubeum* sp. nov. and *Sphaerosporangium cinnabarinum* sp. nov., and transfer of *Streptosporangium viridialbum* Nonomura and Ohara 1960 to *Sphaerosporangium viridialbum* comb. nov. Actinomycetologica 21:11–21.

Ayuso-Sacido, A., and Genilloud, O. 2005. New PCR primers for the screening of NRPS an PKS-I systems in actinomycetes: detection and distribution of these biosynthetic gene sequences inmajor taxonomic groups. Microb. Ecol. 49:10–24.

Berdy, J. 2005. Bioactive microbial metabolites. J. Antibiot. (Tokyo) 58:1–26.

Cao Yan-Ru, Jiang Yi, Xu Li-Hua and Jiang Cheng-Lin: *Sphaerisporangium flaviroseum* sp. nov. and *Sphaerisporangium album* sp. nov., two new species of the genus *Sphaerisporangium* isolated from forest soil in China. Int J Syst Evol Microbiol, 591679–16842009

Chiao, J. S. 2004. [An important mission for microbiologists in the new century-cultivation of the unculturable microorganisms]. Sheng Wu Gong Cheng Xue Bao 20:641–5.

Hou, X. Y. 2001. Collected map of Chinese vegetation. Academic Press, Beijing.

Hayakawa, M., and Nonomura, H.. 1987. Humic acid-vitamin agar, a new medium for the selective isolation of soil actinomycetes. Journal of Fermentation and Bioengineering (Japan) 65:501–509.

Hughes, J. B., Hellmann, J. J., Ricketts, T. H., and Bohannan, B. J. 2001. Counting the uncountable: statistical approaches to estimating microbial diversity. Appl. Environ. Microbiol. 67:4399–406.

Hwang, Y.-B., Lee, M.-Y., Park, H.-J., Han, K., and Kim, E.-S. 2007. Isolation of putative polyene-producing actinomycetes strains via PCR-based genome screening for polyene-specific hydroxylase genes. Process. Biochem. 42:102–107.

Jiang, C. L., and Xu, L. H. 1996. Diversity of soil actinomycetes in Yunnan, China. Appl. Env. Microbiol., 62:244–248.

Jiang, C. L. and Xu, L. H. 1996. Diversity of soil actinomycetes in Yunnan, China. Appl. Env. Microbiol., 62:244–248,1996.

Jiang, C. L., and Xu, L. H. 1997. Microbial Resoucology. Academic Press, Beijing.

Jiang, Y., Duan, S.-R., Tang, S.-K., Cheng, H.-H., Li, W.-J., and Xu, L.-H. 2006. Isolation methods of rare actinomycetes. Microbiology (China). 33:181–183.

Jiang, Y., Wiese, J., Tang, S. K., Xu, L. H., Imhoff, J. F., and Jiang, C. L. 2008. *Actinomycetospora chiangmaiensis* gen. nov., sp. nov., a new member of the family *Pseudonocardiaceae*. Int. J. Syst. Evol. Microbiol. 58:408–13.

Joseph, S. J., Hugenholtz, P., Sangwan, P., Osborne, C. A., and Janssen, P. H. 2003. Laboratory cultivati on of widespread and previously uncultured soil bacteria. Appl. Environ. Microbiol. 69:7210–5.

Metsä-Ketelä, M., Salo, V., Halo, L., Hautala, A., Hakala, J., Mäntsälä, P., and Ylihonko, K. 1999. An efficient approach for screening minimal PKS genes from *Streptomyces*. FEMS Microbiol Lett. 180:1–6.

Pachter, L. 2007. Interpreting the unculturable majority. Nat. Methods. 4:479–80.

Shirling, E. B., and Gottlieb, D. 1966. Methods for characterization of *Streptomyces* species. Int. J. Syst. Bacteriol. 16:313–340.

Wiese, J., Jiang, Y., Tang, S K., Thiel, V., Schmaljohann, R., Xu, L. H., Jiang, C. L., and Imhoff, J. F. 2008. A new member of the family *Micromonosporaceae, Planosporangium flavigriseum* gen.nov.,sp.nov. Int. J. Syst. Evol. Microbiol. 58, 1324–C133.

Mincheol Kim, Hyun-Seok Oh, Sang-Cheol Park and Jongsik Chun. 2014. Towards a taxonomic coherence between average nucleotide identity and 16S rRNA gene sequence similarity for species demarcation of prokaryotes. Int. J. Syst. Evol. Microbiol. 64, 346–351.

Xu, P., Li, W. J., Xu, L. H., and Jiang, C. L. 2003. A microwave based method for genomic DNA extraction from actinomycetes. Microbiology (China). 30:82–84.

Yang, Y., Jiang, Y., Yim, M., Xu, P., Lu, T., and Xu, L. H. 2007. Synthesis genomic of secondary metabolites of actinomycetes. Microbiology (China). 27:68–71.

Zengler, K., Toledo, G., Rappe, M., Elkins, J., Mathur, E. J., Short, J. M., and Keller, M. 2002. Cultivating the uncultured. Proc. Natl. Acad. Sci. U S A 99:15681–6.

